# Dephosphorylation of the NPR2 guanylyl cyclase contributes to inhibition of bone growth by fibroblast growth factor

**DOI:** 10.1101/193847

**Authors:** Leia C. Shuhaibar, Jerid W. Robinson, Ninna P. Shuhaibar, Jeremy R. Egbert, Giulia Vigone, Valentina Baena, Deborah Kaback, Siu-Pok Yee, Robert Feil, Melanie C. Fisher, Caroline N. Dealy, Lincoln R. Potter, Laurinda A. Jaffe

**Author notes:** For correspondence (LCS); (LRP); (LAJ).

## Abstract

Activating mutations in fibroblast growth factor (FGF) receptor 3 and inactivating mutations in the NPR2 guanylyl cyclase cause similar forms of dwarfism, but how these two signaling systems interact to regulate bone growth is poorly understood. Here, by use of a mouse model in which NPR2 cannot be dephosphorylated, we show that bone elongation is opposed when NPR2 is dephosphorylated and thus produces less cyclic GMP. By developing an in vivo imaging system to measure cyclic GMP levels in intact tibia, we show that FGF-induced dephosphorylation of NPR2 decreases its guanylyl cyclase activity in growth plate chondrocytes in living bone. Thus FGF signaling lowers cyclic GMP in the growth plate, which counteracts bone elongation. These results define a new component of the signaling network by which activating mutations in the FGF receptor inhibit bone growth.

## Introduction

Longitudinal growth of limbs and vertebrae depends on division and differentiation of chondrocytes within the cartilage growth plate, resulting in formation of a scaffold that is subsequently mineralized (***Kozhemyakina et al., 2015***). These processes are tightly controlled by multiple regulatory pathways. One such regulator is the natriuretic peptide-stimulated guanylyl cyclase, natriuretic peptide receptor 2 (NPR2), also known as guanylyl cyclase B, which is found in growth plate chondrocytes and promotes bone elongation (***Yasoda et al., 1998; Yamashita et al., 2000; Chuso et al., 2001; Tamura et al., 2004; Bartels et al., 2004***). Inactivating mutations in NPR2, which reduce cGMP, result in shorter bones, and cause acromesomelic dysplasia Maroteaux type dwarfism (***Tamura et al., 2004; Bartels et al., 2004; Khan et al., 2012; Geister et al., 2013; Nakao et al., 2015***). Conversely, activating mutations of NPR2 result in longer bones (***Miura et al., 2012; Hannema et al., 2013; Miura et al., 2014***). In addition to its role in regulating bone length, cGMP also regulates bone density (***Kondo et al., 2012***; ***Kalyanaraman et al., 2017***).

Activation of the guanylyl cyclase activity of NPR2 requires the extracellular binding of C-type natriuretic peptide (CNP) and also the phosphorylation of multiple intracellular juxtamembrane serines and threonines (***Potter, 1998; Potter, 2011*W**) (***Figure 1A***). If these sites are mutated to alanine, CNP-dependent guanylyl cyclase activity is reduced to only 6% of that of the wild type protein (***Potter and Hunter, 1998***). Correspondingly, if the 7 serines and threonines are mutated to the phosphomimetic amino acid glutamate (NPR2-7E), the CNP-dependent guanylyl cyclase activity is the same as that of the wild type phosphorylated enzyme, but does not decrease in response to stimuli that dephosphorylate and inactivate the wild type protein (***Yoder et al., 2012; Shuhaibar et al., 2016***) (***Figure 1A***). Our previous analysis of mice in which both copies of *Npr2* were globally replaced with a sequence encoding NPR2-7E (*Npr2*^*7E/7E*^) demonstrated that dephosphorylation of NPR2 is a physiological mediator of hormonal signaling in ovarian follicles (***Shuhaibar et al., 2016***). These findings led us to investigate whether the phosphorylation of NPR2 could also be a mediator of growth factor signaling in bones.

**Figure 1.**
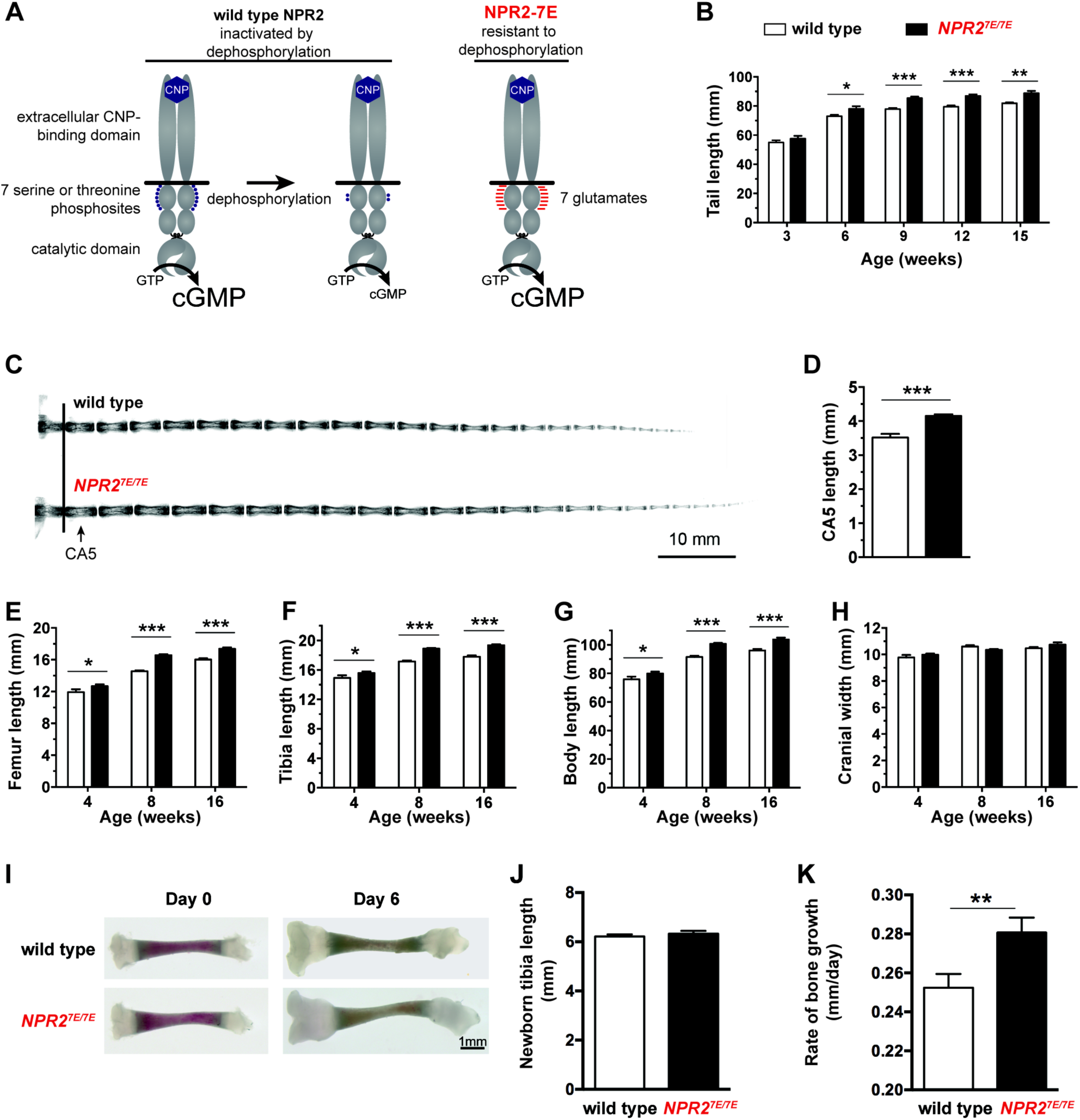
Mutation of the 7 regulatory serines and threonines of NPR2 to glutamates increases bone length. (**A**) Regulation of NPR2 activity by the phosphorylation state of the 7 juxtamembrane serines and threonines on each NPR2 monomer. Purple dots represent phosphates on these serines and threonines. Red lines represent glutamates that are substituted for the serines and threonines. (**B**) *Npr2*^*7E/7E*^ mice have longer tails. Measurements were made from 17-19 live mice of each genotype. (**C**) X-rays of representative tails from mice euthanized at 18 weeks of age. (**D**) Increased CA5 vertebra length in *Npr2*^*7E/7E*^ mice compared with wild type, determined from x-ray measurements of tails at 18 weeks of age (10-13 mice of each genotype). (**E,F**) Longer femurs (**E**) and tibias (**F**) in *Npr2*^*7E/7E*^ vs wild type mice. (**G**) Increased body length in *Npr2*^*7E/7E*^ vs wild type mice. (**H**) No difference in cranial width comparing *Npr2*^*7E/7E*^ and wild type mice. For **E-H**, each bar indicates measurements from 10-31 mice that were euthanized at the indicated ages. (**I - K**) Growth of tibia from newborn mice, cultured in vitro for 6 days in the presence of 1 µM CNP. (**I**) Representative tibia before and after 6 days of culture. (**J**) Tibia length immediately after dissection.(**K**) Rates of longitudinal bone growth. For (**J**) and (**K**), 28 wild type and 30 *Npr2*^*7E/7E*^ bones were measured. All graphs show mean ± s.e.m. Data were analyzed by unpaired t-tests, with the Holm-Sidak correction for multiple comparisons where appropriate. T-tests rather than ANOVA were used because we were only interested in comparisons between genotypes at a given age, rather than comparisons across ages. * = p ≤ 0.05; ** = p ≤ 0.01; *** = p ≤ 0.001.

Another essential regulator of bone elongation is FGF receptor 3 (FGFR3); activating mutations of FGFR3 inhibit bone growth, causing achondroplasia and other forms of dwarfism (***Rousseau et al., 1994; Shiang et al., 1994; Naski et al., 1998; Wang et al., 1999; Lorget et al., 2012; Lee et al., 2017; Ornitz and Legeai-Mallet, 2017***). Conversely, mice lacking FGFR3 have longer bones (***Deng et al., 1996***). Intriguingly, the reduced bone length seen with activating mutations of FGFR3 resembles that seen with inactivating mutations of NPR2, and activation of NPR2 by increasing CNP opposes the decrease in bone growth caused by activating mutations of FGFR3 (***Yasoda et al., 2004; Yasoda et al., 2009; Lorget et al., 2012; Wendt et al., 2015***). Likewise, the increased bone length seen with mice lacking FGFR3 resembles that seen with activating mutations of NPR2 (***Miura et al., 2012; Hannema et al., 2013; Miura et al., 2014***). Furthermore, studies of fibroblasts and chondrogenic cells derived from embryonic carcinomas and chondrosarcomas have indicated that FGF signaling decreases NPR2 activity (***Chrisman and Garbers, 1999; Ozasa et al., 2005; Robinson et al., 2017),*** and that the inactivation is due to dephosphorylation of NPR2 (***Robinson et al., 2017***). These studies suggested to us that dephosphorylation and inactivation of NPR2 could be a mechanism by which FGF decreases elongation of bones in vivo.

Here we tested this hypothesis by investigating the growth of bones in *Npr2*^*7E/7E*^ mice in which NPR2 cannot be inactivated by dephosphorylation, and found that their bones are longer than those of wild type mice. We then developed a live tissue imaging system for monitoring the guanylyl cyclase activity of NPR2 in intact growth plates from wild type and *Npr2*^*7E/7E*^ mice that express a FRET sensor for cGMP. Using this system, we found that FGF-induced dephosphorylation and inactivation of NPR2 decreases cGMP in growth plate chondrocytes, thus contributing to FGF-dependent decreases in bone growth.

## Results

### Mice in which NPR2 cannot be dephosphorylated have longer bones

*Npr2*^*7E/7E*^ mice, in which NPR2 is modified to mimic a constitutively phosphorylated protein (***Shuhaibar et al., 2016***) (***Figure 1A***), have longer bones. Measurements at 8-16 weeks of age showed that the tails, femurs, tibias, and body of *Npr2*^*7E/7E*^ mice were 8 to 14% longer than wild type (***Figure 1B,E,F,G***). Differences in limb and body length were seen as early as 4 weeks (***Figure 1E-G***). The fifth caudal vertebra was 17% longer, as measured at 18 weeks (***Figure 1C,D***). These results indicate that when NPR2 cannot be inactivated by dephosphorylation, endochondral bone growth is increased. The width of the cranium, which is dependent on membranous, not endochondral, bone growth, was unaffected (***Figure 1H***). The measured increases in body and bone length, which resulted from an inability of NPR2 to be inactivated by dephosphorylation, were comparable in magnitude to those previously reported for mice in which NPR2 activity was increased by overexpression of the NPR2 agonist CNP (***Yasoda et al., 2004; Yasoda et al., 2009***).

### *Npr2*^*7E/7E*^ mice have longer bones due to an intrinsic effect on bone

Because NPR2 is expressed in tissues outside of the bone growth plate, including the pituitary and gastro-intestinal tract (***Tamura and Garbers, 2003; Sogawa et al., 2010***), we tested whether the longer bones of the *Npr2*^*7E/7E*^ mice resulted from a direct action on bone rather than an indirect action on other tissues. To do this, we cultured isolated tibia from 3-4 day old mice, such that we could measure growth rates in the absence of extrinsic hormonal or nutritional effects.

When first dissected, *Npr2*^*7E/7E*^ and wild type tibia did not differ in length (**Figure 1 I, J**). The absence of a detectable difference in length is consistent with previous studies showing no difference in body length at birth comparing mice with inactivated NPR2 and wild type mice (***Tamura et al., 2004***). Likewise, little or no difference in body or bone length is seen at birth of humans with inactivating NPR2 mutations (***Bartels et al., 2004***). These previous studies indicate that NPR2 activity is not required for prenatal bone elongation. However, when bones of newborn mice are placed in culture in the presence of CNP, bone elongation over a 6-day period is reduced if the *Npr2* gene is disrupted (***Tamura et al., 2004***). Correspondingly, when cultured in the presence of 1 µM CNP and measured over a period of 6 days, tibia from *Npr2*^*7E/7E*^ mice showed a 12% larger rate of elongation compared with wild type tibia (***Figure 1 I, K***). These results show that *Npr2*^*7E/7E*^ mice have longer bones due to an intrinsic effect on bone.

### Cyclic GMP measurements in live growth plates

To establish a method to investigate the signaling mechanisms underlying the increased bone growth in *Npr2*^*7E/7E*^ mice, we developed a live imaging technique for monitoring NPR2 guanylyl cyclase activity in intact growth plates, using mice that globally express a sensor for cGMP, cGi500 (***Russwurm et al., 2007; Thunemann et al., 2013; Shuhaibar et al., 2015***). NPR2 activity has been previously measured using homogenates of fetal tibia (***Yasoda et al., 1998***) and membrane preparations from chondrocyte cell lines (***Osaza et al., 2005; Robinson et al., 2017***). The availability of mice expressing a sensor for cGMP, combined with the development of a new method to visualize chondrocytes in live and intact growth plates by confocal microscopy, allowed us to measure NPR2 activity under physiological conditions and with high spatial and temporal resolution.

cGi500 is comprised of cyan fluorescent protein (CFP) and yellow fluorescent protein (YFP), linked by a cGMP-binding domain; the sensitivity of cGi500 to cGMP is based on Förster resonance energy transfer (FRET). Binding of cGMP to the linker domain causes the CFP and YFP domains to move farther apart. The proximity of CFP and YFP can be detected by imaging cells using a 440 nm laser to excite CFP; the energy emitted by CFP then excites YFP in proportion to the distance between the 2 fluorophores. Thus measurement of the relative intensities of the light emitted by CFP and YFP provides an indicator of the cGMP concentration. The CFP/YFP emission ratio is higher when the cGMP concentration is higher, with a half-maximal response at 500 nM cGMP (**Russwurm et al., 2007**).

Tibias were removed from newborn mice expressing cGi500, and the growth plate was exposed by cutting away the overlying tissue (***Figure 2A***), thus allowing imaging and rapid exchange of solutions. A tibia was then placed in a perfusion slide (***Figure 2B,C***) on the stage of a confocal microscope. Excitation of CFP resulted in fluorescence in all regions of the growth plate (***Figure 2D***), allowing identification of chondrocytes at different stages of development, based on the shape of the cells: round (also called resting), columnar (also called proliferating), and hypertrophic (***Kozhemyakina et al., 2015; Ornitz and Legeai-Mallet, 2017***). Columnar cells that are beginning to increase their volume, referred to as “prehypertrophic”, express both NPR2 and FGFR3 (***Yamashita et al., 2000; Chusho et al., 2001; de Frutos et al., 2007; Kozhemyakina et al., 2015)***. We used this columnar/prehypertrophic region for the FRET measurements (***Figure 2E***), such that FGF effects on NPR2 activity could be investigated.

**Figure 2.**
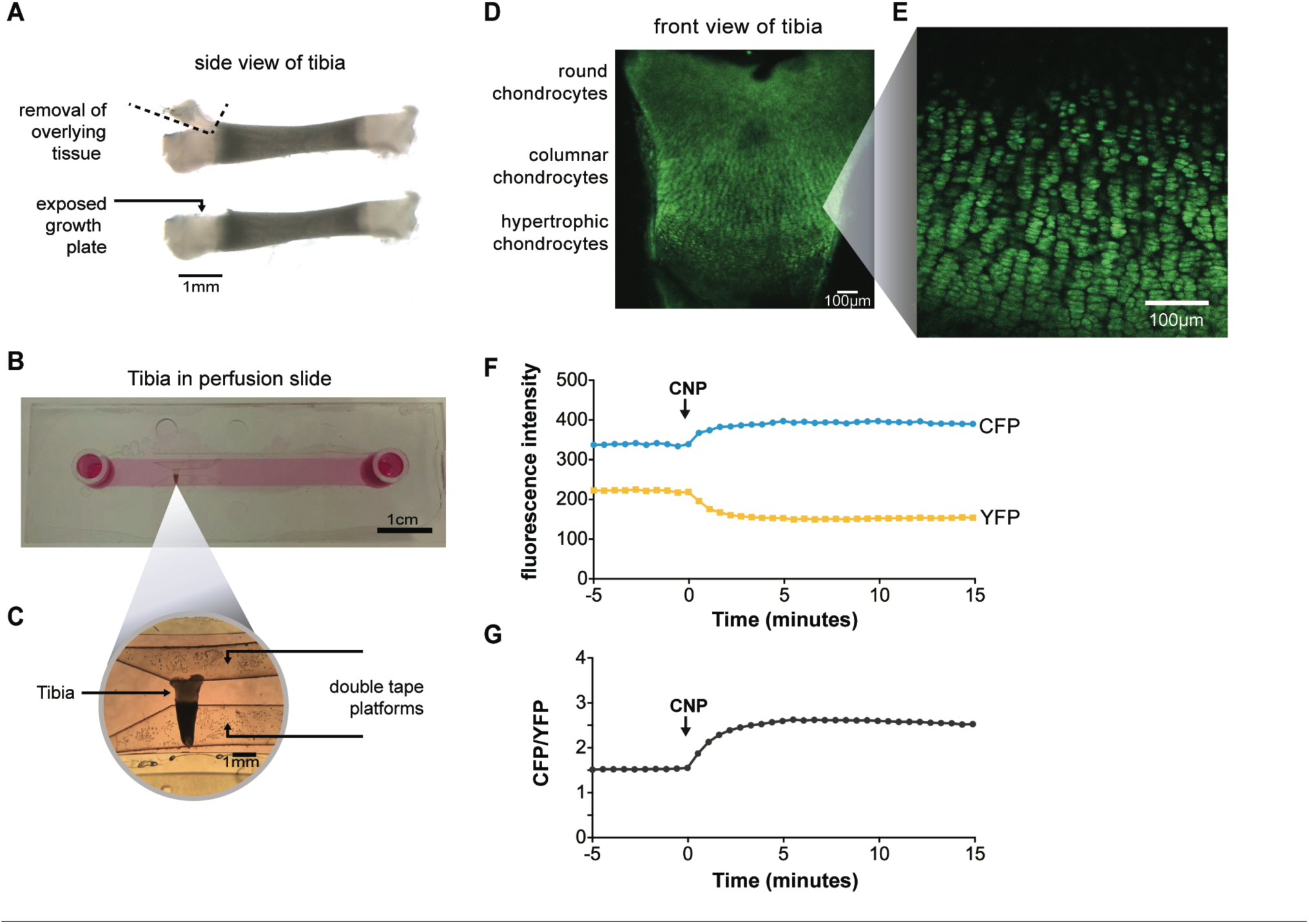
Imaging of CNP-stimulated cGMP generation in chondrocytes within intact growth plates.(**A**) Preparation of tibia from a newborn mouse expressing the cGi500 sensor for cGMP. (**B**) Tibia in a perfusion slide in which medium could be flowed across the growth plate surface while imaging.(C) Higher magnification image of the tibia in the perfusion slide. (**D**) Confocal image of fluorescence of the entire growth plate. (**E**) Higher magnification image of the columnar/prehypertrophic region that was used for measurements of cGMP (from a different tibia from that shown in D). (**F,G**) Time courses of CFP and YFP emission intensities and the CFP/YFP ratio, before and after perfusion of 100 nM CNP.

To measure the CNP-dependent guanylyl cyclase activity of NPR2, we first recorded baseline images of CFP and YFP emission in the absence of CNP. After establishing the baseline ratio, we perfused 100 nM CNP through the imaging chamber. In response to the CNP, the CFP emission intensity increased and the YFP emission intensity decreased (***Figure 2F***), resulting in a higher CFP/YFP ratio (***Figure 2G***) and indicating an increase in cGMP due to activation of NPR2. Thus, confocal microscopy of isolated tibia expressing cGi500 allowed us to monitor NPR2 activity in the live and intact growth plate for the first time.

### FGF reduces NPR2 guanylyl cyclase activity in the growth plate

To investigate whether FGF signaling reduces NPR2 activity within growth plate chondrocytes, we incubated cGi500-expressing tibias with or without FGF18, and imaged the fluorescence of chondrocytes in the columnar/prehypertrophic region. Among the many FGF isoforms, we used FGF18 because it is one of the two redundant FGF isoforms that function to activate FGFR3 in the growth plate (***Ornitz and Marie, 2015; Hung et al., 2016***). In tibias without FGF18 pretreatment, perfusion of CNP through the imaging chamber caused a large increase in the concentration of cGMP (***Figure 3A***). However, if the tibias were pretreated with FGF18, the CNP-induced increase in cGMP was smaller (***Figure 3A,G***). The baseline cGMP level before CNP perfusion was the same with and without FGF pretreatment (***Figure 3A***). These results indicate that in chondrocytes within a living growth plate, FGF signaling reduces the activity of the NPR2 guanylyl cyclase. Similar results were obtained with FGF18 or control pretreatment times of 80-140 minutes (***Figure 3A,G***), or 30-50 minutes (***Figure 3 -- figure supplement 1***).

**Figure 3.**
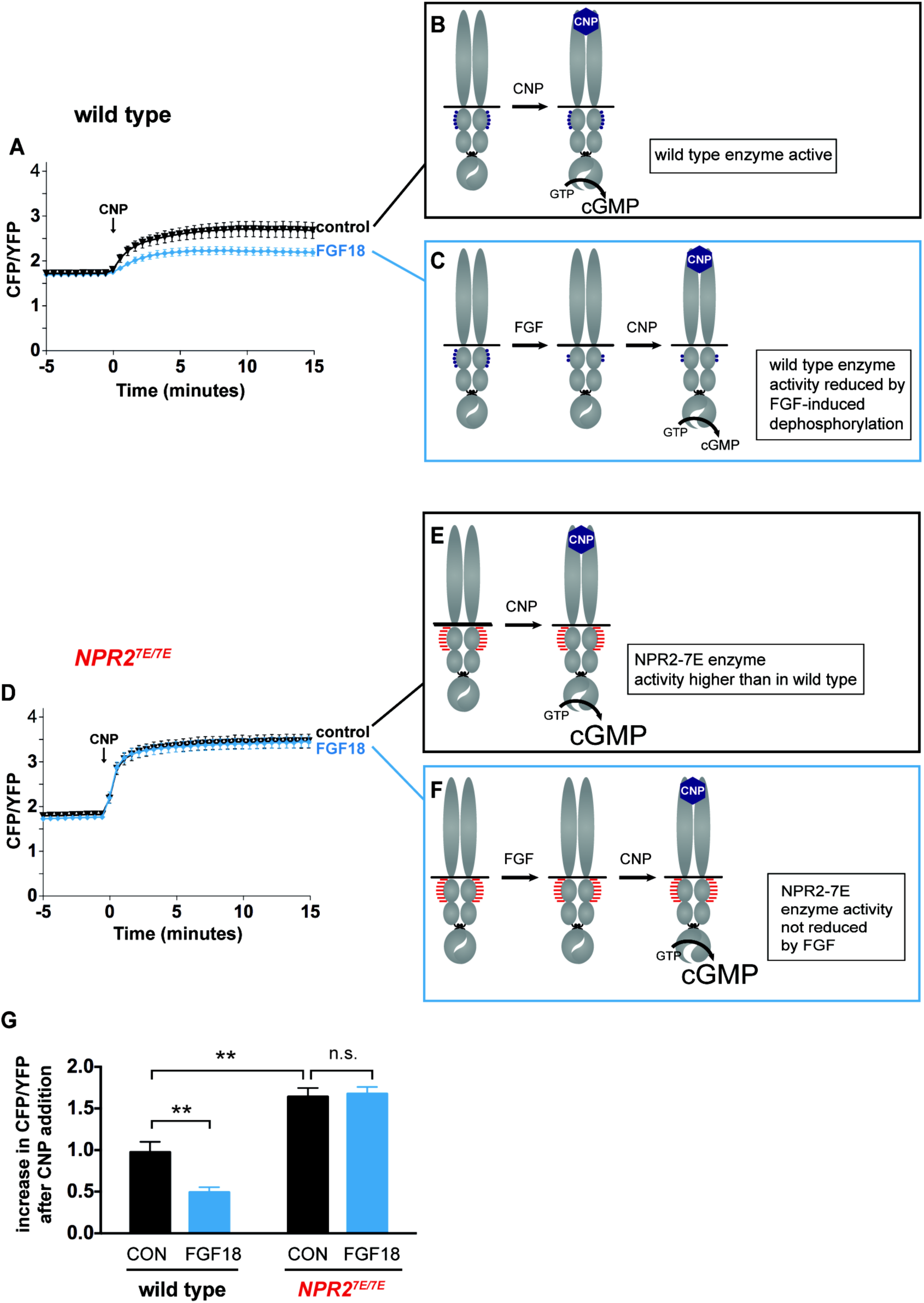
FGF reduces NPR2 guanylyl cyclase activity in growth plate chondrocytes from wild type mice, but not in those from *Npr2*^*7E/7E*^ mice. (**A**) Time course of CFP/YFP intensity ratios after 100 nM CNP was perfused across a wild type growth plate, comparing tibias that were preincubated for 80-140 minutes with a control solution (1 μg/ml heparin) (n = 11) or with FGF18 (0.5 µg/ml FGF18 + 1 μg/ml heparin) (n = 14). CNP was present for the remainder of the recording period. An increase in the CFP/YFP ratio in response to CNP indicates cGMP production by NPR2. This and other graphs show mean ± s.e.m. (**B,C**) Schematic diagrams depicting the dephosphorylation of NPR2 in response to FGF, as demonstrated in rat chondrosarcoma cells (***Robinson et al., 2017***), and the effect on NPR2 guanylyl cyclase activity, in wild type tibia. Without FGF treatment, the 7 regulatory serines and threonines on each monomer are mostly phosphorylated (depicted by 5 purple dots).After FGF treatment, the regulatory sites are mostly dephosphorylated (depicted by 2 purple dots).(D) Time course of CFP/YFP intensity ratios after 100 nM CNP was perfused across an *Npr2*^*7E/7E*^growth plate, comparing tibias that were preincubated for 80-140 minutes with a control solution (1μg/ml heparin) (n = 10) or with FGF18 (0.5 µg/ml FGF18 + 1 μg/ml heparin) (n = 11). (**E,F**) Schematic diagrams depicting NPR2 with 7 glutamates on each monomer (indicated by 7 red lines) and the lack of effect of FGF on NPR2 enzyme activity. (**G**) Increases in CFP/YFP ratios after CNP perfusion, comparing data from wild type tibias with and without FGF pretreatment (from **A**) and data from *Npr2*^*7E/7E*^ tibias with and without FGF pretreatment (from **D**). Values were determined by subtracting the mean CFP/YFP ratio 0-5 minutes before CNP perfusion from the mean ratio 10-15 minutes after CNP perfusion. Data were analyzed by unpaired t-tests, with the Holm-Sidak correction for multiple comparisons. T-tests rather than ANOVA were used because we were only interested in a subset of comparisons that tested specific hypotheses. ** = p ≤ 0.01.

### The FGF-induced decrease in guanylyl cyclase activity is caused by NPR2 dephosphorylation

To test whether FGF reduces the guanylyl cyclase activity of NPR2 in growth plate chondrocytes by dephosphorylating NPR2, as diagrammed in ***Figure 3B,C***, we examined cGMP production in growth plates from tibias of *Npr2*^*7E/7E*^ mice, such that FGF could not dephosphorylate NPR2. As with wild type, growth plates from *Npr2*^*7E/7E*^ mice showed an increase in cGMP when exposed to CNP (***Figure 3D***). In fact, the CNP-stimulated cGMP increase was greater for *Npr2*^*7E/7E*^ vs wild type (***Figure 3G***), as would be expected if NPR2 in the wild type growth plate was partially dephosphorylated as a result of signaling by endogenous hormones or growth factors (compare diagrams in ***Figures 3B and E***). Importantly, in contrast to wild type, growth plates from *Npr2*^*7E/7E*^ mice showed a similar CNP-stimulated increase in cGMP with or without FGF pretreatment (***Figure 3D, G***), consistent with the diagram in ***Figure 3F***. These results indicate that FGF signaling reduces cGMP production by dephosphorylating NPR2 in the growth plate.

Under the culture conditions used for the FRET measurements, a difference in cGMP levels in growth plate chondrocytes of *Npr2*^*7E/7E*^ and wild type newborn mice was seen only after the addition of CNP to the medium (***Figures 3A,D,G***). The concentrations of CNP in the extracellular space around chondrocytes of tibia in vivo are not known exactly, but are evidently sufficient to activate NPR2, since mutations that inactivate NPR2 inhibit bone elongation (see Introduction). Thus, we can conclude that *Npr2*^*7E/7E*^ chondrocytes in vivo have higher cGMP concentrations compared to wild type chondrocytes.

## Discussion

Our findings identify two new components of the signaling network by which FGFR3 and NPR2 signaling regulate bone growth. Firstly, our results indicate that phosphorylation of the NPR2 guanylyl cyclase promotes bone elongation; this conclusion is based on our finding that a mutation that prevents NPR2 dephosphorylation results in longer bones. Secondly, our results indicate that part of the mechanism by which FGF signaling inhibits bone elongation is by decreasing NPR2 phosphorylation and activity. However, mice lacking the FGFR3 receptor (***Deng et al., 1996***) show a greater increase in bone length than we report here for mice with the mutation that prevents dephosphorylation of NPR2, consistent with previous studies identifying multiple mechanisms by which FGF signaling opposes bone elongation (***Ornitz and Legeai-Mallet, 2017***).

The working model in ***Figure 4*** shows our findings in the context of other knowledge about the cross-talk between FGFR3 and NPR2 signaling. The major pathway by which FGFR3 signaling decreases bone elongation is by activation of MAP kinase (see Ornitz and Legeai-Mallet, 2017). The major pathway by which NPR2 guanylyl cyclase activity increases bone elongation is by elevating cGMP and activating the cGMP-dependent protein kinase PRKG2 (***Pfeifer et al.,1996; Chikuda et al., 2004***). Previous studies have established that application of CNP to increase NPR2 guanylyl cyclase activity in chondrocytes, thus elevating cGMP and stimulating cGMP-dependent protein kinase, inhibits FGF-induced MAP kinase activation (***Yasoda et al., 2004; Krejci et al., 2005; Ozasa et al., 2005; Kamemura et al., 2017***), which would counteract FGF-inhibition of bone growth (solid blue line in ***Figure 4***). Other mechanisms (for example, see ***Kawasaki et al., 2008***) may also contribute to how PRKG2 activity increases bone growth (dotted blue line in ***Figure 4***).

**Figure 4.**
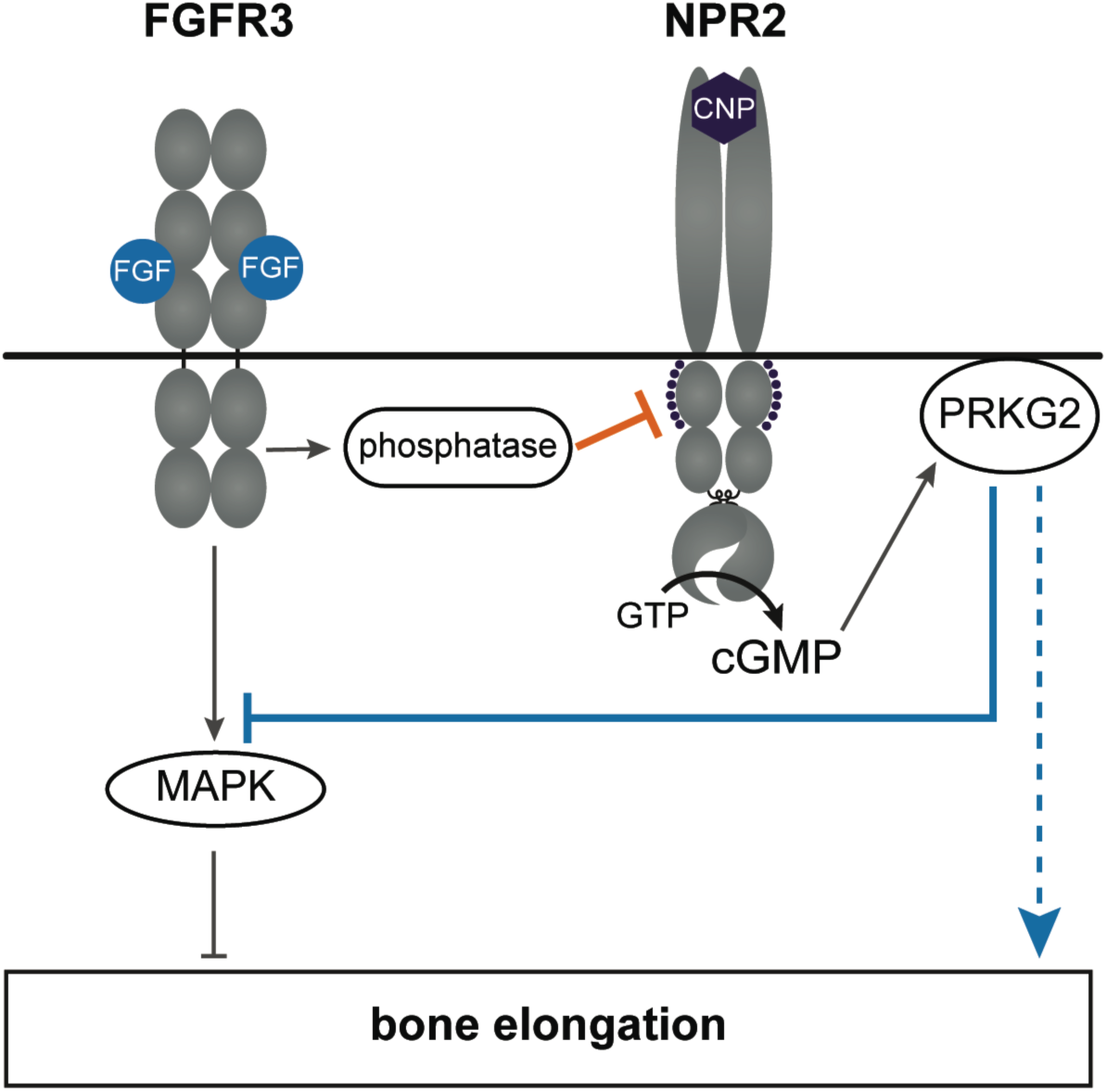
A working model of the crosstalk between NPR2 and FGFR3 signaling pathways in regulating bone elongation, emphasizing the role of the phosphorylation state of NPR2. See main text for discussion. This diagram shows only one aspect of FGFR3 signaling in the growth plate; see Karuppaiah et al. (2016) and Ornitz and Legeai-Mallet (2017) for current models showing other components as well.

Our present findings identify another level of crosstalk between FGFR3 and NPR2 signaling, indicated by the orange line in ***Figure 4***. By causing dephosphorylation of NPR2, FGF signaling lowers cGMP, thus opposing bone elongation. How FGF signaling dephosphorylates NPR2 remains to be determined; this could occur by inactivation of the kinase(s) that phosphorylate NPR2, and/or by activation of the phosphatase(s) that dephosphorylate it, neither of which has been identified. Activation of the protein phosphatase PP2A is one possibility, since PP2A is activated by FGF signaling in the RCS chondrocyte cell line (***Kolupaeva et al., 2013***), and purified PP2A can dephosphorylate NPR2 in vitro (***Potter, 1998***).

Other groups have used fluorescence microscopy to quantify chondrocyte movement, division, and volume in live avian growth plate cartilage (***Li et al., 2015***), and to measure chondrocyte density in fixed mouse mandibular cartilage (***He et al., 2017***). However, the methods that we have developed for imaging the growth plate of mammalian bones are unique in that they allow rapid manipulation of the chemical environment surrounding the growth plate and real-time measurements of changes in signaling pathways within the intact tissue. These methods are broadly applicable to studies of signaling by other hormones and growth factors that might affect cGMP levels in growth plate chondrocytes. With mice expressing related FRET sensors for cAMP (***Calebiro et al., 2009***), these methods could also be readily applied to studies of signaling by hormones such as parathyroid hormone related protein that regulate chondrocyte and osteoblast growth and differentiation by way of cAMP (***Chagin et al., 2014; Kozhemyakina et al., 2015; Esbrit et al., 2016***).

Ongoing clinical trials have shown that in patients with achondroplasia, in which skeletal dwarfism is caused by a constitutively active form of FGFR3, stimulation of NPR2 by subcutaneous injection of a hydrolysis-resistant analog of CNP increases bone length (***Klag and Horton, 2016***). These results are consistent with the increased bone growth that results from increasing CNP in mouse models of achondroplasia (***Yasoda et al., 2004; Yasoda et al., 2009; Lorget et al., 2012; Wendt et al., 2015***). Studies using a mouse model overexpressing a constitutively active form of FGFR3 showed that applying CNP or a CNP analog in vivo or in vitro completely rescued the growth defect (***Yasoda et al., 2004; Yasoda et al., 2009; Wendt et al., 2015***). However, in a study involving mice in which one allele of the *Fgfr3* gene was replaced with a constitutively active form, as occurs in achondroplasia, it was found that treatment with the CNP analog increased the growth rate of cultured tibia only partially, from ∼40% to ∼70% of wild type (***Lorget et al., 2012***). Such incomplete rescue might be expected, considering that NPR2 guanylyl cyclase activity depends not only on the presence of CNP but also on phosphorylation of multiple regulatory serines and threonines of NPR2 (***Figure 1A***). CNP levels are elevated in patients with achondroplasia, suggesting that NPR2 in their chondrocytes is resistant to CNP (***Olney et al., 2015***), which could be due to NPR2 dephosphorylation. Thus, if NPR2 phosphorylation could be increased, either by activating the kinase that phosphorylates NPR2 or by inhibiting the phosphatase that dephosphorylates NPR2, these processes could potentially be targeted to enhance the therapeutic stimulation of NPR2 activity by CNP as a treatment for achondroplasia.

## Materials and Methods

### Mice

Two mouse lines were used for this study: *Npr2*^*7E/7E*^ (***Shuhaibar et al., 2016***), and cGi500 (***Thunemann et al., 2013***). All experiments were conducted as approved by the University of Connecticut Health Center and the University of Minnesota animal care committees.

### Measurements of tail, bone, and body lengths

Tail lengths of live mice were measured at 3 week intervals. The lengths of the fifth caudal vertebrae were measured from euthanized mice, using a Faxitron cabinet x-ray system. Body lengths were measured from the tip of the nose to the base of the tail, using euthanized mice and a digital caliper. Femur and tibia lengths and cranial width were measured using excised bones and a digital caliper. For all measurements, there were approximately equal numbers of males and females of each genotype.

### Measurement of tibia growth in vitro

Tibias were dissected from 3-4 day old mice, and cultured as previously described (***Tamura et al., 2004***) on Millicell organotypic membranes (PICMORG50; Merck Millipore Ltd, Cork, IRL), in BGJb medium (Fitton-Jackson modification) (Life Technologies, Grand Island, NY) with 0.1% BSA, 100 units/ml each of penicillin and streptomycin, and 1 µM CNP (Phoenix Pharmaceuticals, Burlingame, CA). The medium was changed every 2 days. Tibias were photographed using a Leica stereoscope, for measurement of length before and after a 6-day culture period.

### Measurement of cGMP

Cyclic GMP was measured using tibias dissected from newborn mice (day 0-1) that globally expressed one or two copies of the cGi500 FRET sensor inserted into the *Rosa26* locus (***Thunemann et al., 2013***). Isolated tibias were placed on Millicell membranes, in the medium described above, and used for imaging over the next 2 days. Prior to imaging, the epiphysis region of the proximal end of the tibia was slit with a razor blade to expose the growth plate (***Figure 2A***). This was accomplished by placing the tibia, dorsal side up, in a 500 μm deep channel in a plastic slide (ibidi USA, Fitchburg, WI; cat. no. 80176, special order with no adhesive). The depth of the channel was modified from 400 to 500 μm deep by adding a piece of tape on each side of the channel. The epiphysis region was then slid through the edge of a razor blade on the slide surface. This procedure resulted in a 500 μm thick piece of tissue underlying the growth plate surface. The overlying flap of tissue was trimmed away, and the tibia was placed in a 4-well plate (Nunc 176740; Thermo Scientific, Rochester, NY) containing 0.5-1 ml of medium/well at 37^o^C with 5% CO_2_, on a rotating platform. Where indicated, FGF18 (PeproTech, Rocky Hill, NJ) was added at a saturating concentration (0.5 μg/ml), along with 1 μg/ml heparin (Sigma-Aldrich, H4784, St. Louis, MO).Heparin was included because it enhances FGF receptor activation (***Ornitz and Marie, 2015***). Control samples were incubated with heparin only.

After the indicated incubation time, the distal half of the tibia was cut off, and the proximal piece placed, cut edge up, in a 600 μm deep channel in a plastic slide with access wells for perfusion (ibidi USA; cat. no. 80186; special order with no adhesive) (***Figure 2B***). Silicon grease was applied around the rim of the channel. A coverslip was prepared by adhering to its surface two 200 µm thick pads that were separated by a 1-1.5 mm space. The pads were made from 2 layers of Scotch double sided tape, each 100 μm thick. The coverslip was then placed over the tibia, such that the bone spanned the two pads and the surface of the growth plate was separated by ∼200 μm from the surface of the coverslip (***Figure 2C***). The coverslip was pressed down gently against the silicon grease on the perfusion slide, such that the uncut surface of the tibia was held against the perfusion slide. The slide was then inverted and filled with media, by way of ports on the slide. This resulted in an assembly in which media could be flowed under the bridge formed by the tibia resting on the tape pads. For each experiment, the separation of the growth plate surface from the coverslip was confirmed by measuring the distance between these surfaces using the confocal microscope software.

The growth plate was imaged using a Pascal confocal system (Carl Zeiss Microscopy, Thornwood, NY). Imaging was performed at ∼34^o^C. The whole growth plate was imaged with a 10x/0.45 N.A. water immersion objective (***Figure 2D***). For FRET measurements, columnar/prehypertrophic chondrocytes were imaged using a 25x/0.8 NA water immersion objective (***Figure 2E***). To avoid evaporation during time-lapse imaging, Immersol (Carl Zeiss Microscopy) was used to form the contact with the coverslip. The pinhole was set at maximum, resulting in an optical section thickness of ∼24 μm. FRET measurements were performed and analyzed as previously described (***Norris et al., 2009; Shuhaibar et al., 2015***). Images were collected at zoom 0.7, using 3.2 second scans at 30 second intervals, for 5 minutes before CNP perfusion and 15 minutes afterwards. Files were saved as 12-bit images. The fluorescence signal was 2 – 6 times the background level. Measurements were corrected for background and for spectral bleedthrough of light emitted by CFP into the YFP channel. Graphs show measurements from 490 x 490 µm regions.

## Acknowledgments

We thank Tracy Uliasz for technical assistance, Liping Wang and David Rowe for help with x-ray imaging, and Melina Schuh for critical reading of the manuscript. This work was supported by grants from the National Institutes of Health (R37HD014939 to LAJ, R01GM098309 to LRP, T32DK007203 to JWR, and R90DE022526 to NPS), and from the Fund for Science (to LCS, CND, LRP, and LAJ).

---

**Figure 3 - figure supplement 1.**
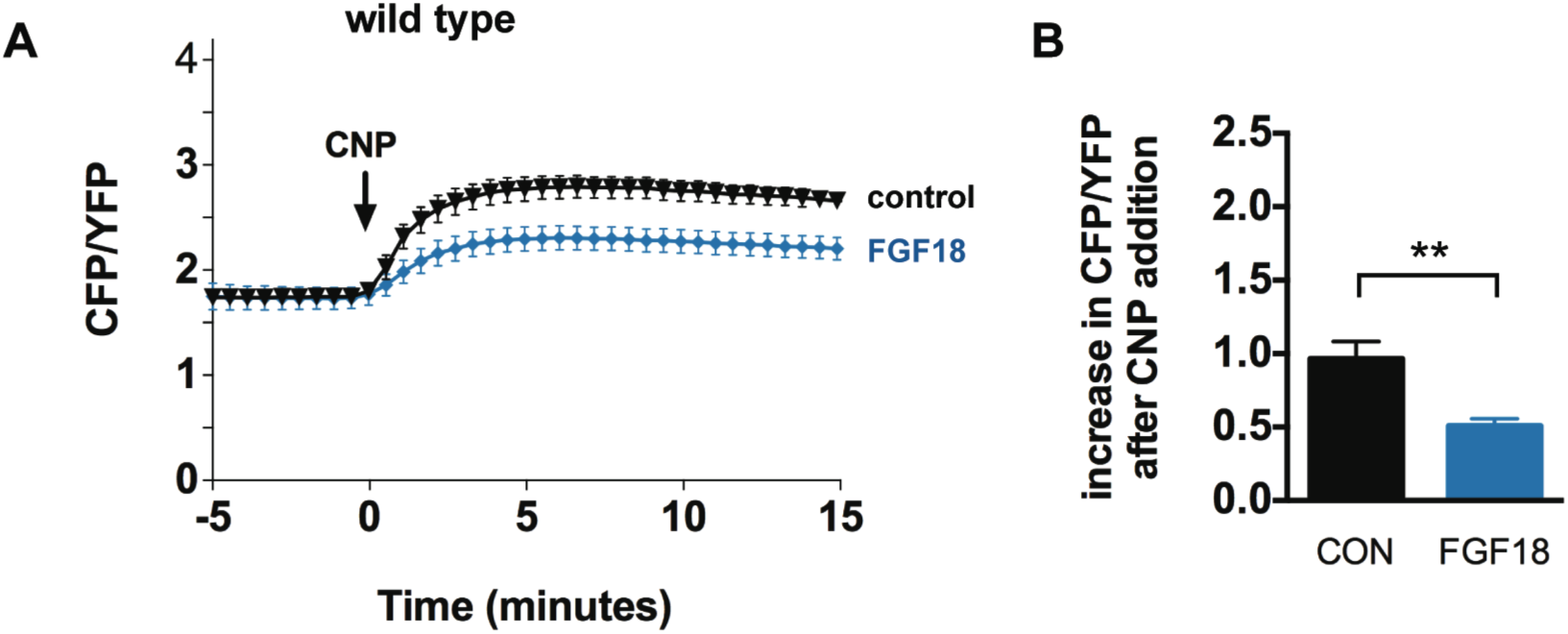
Inhibition of NPR2 activity by a 30-50 minute preincubation with FGF. (**A**) Time course of CFP/YFP intensity ratios after 100 nM CNP was perfused across a wild type growth plate, comparing tibias that were preincubated for 30-50 minutes with a control solution (1 μg/ml heparin) (n = 8) or with FGF18 (0.5 µg/ml FGF18 + 1 μg/ml heparin) (n = 8). (**B**) Increases in CFP/YFP ratios after CNP perfusion, comparing data from wild type tibias with and without FGF pretreatment (from **A**). Values were determined by subtracting the mean CFP/YFP ratio 0-5 minutes before CNP perfusion from the mean ratio 10-15 minutes after CNP perfusion. Data were analyzed by unpaired t-tests. ** = p ≤ 0.01.

